# Cross-Sectional Physiological and Neuromuscular Profiling of Elite and Recreational University Badminton Athletes: Preliminary Benchmarks for Exercise-Based Injury Risk Stratification

**DOI:** 10.64898/2026.06.02.729234

**Authors:** Humayon Ahmed, Md. Moznuzzaman, Md. Khalid Hasan, Jishan Ahammad Shohag, Mehedi Hasan, Asif Abdullah, Farjana Akter Boby

**Affiliations:** Department of Physical Education and Sports Science, Jashore University of Science and Technology, Jashore, Bangladesh; Department of Electrical and Electronic Engineering, Jashore University of Science and Technology, Jashore, Bangladesh; Department of Biomedical Engineering, Jashore University of Science and Technology; Department of Physical Education and Sports Science, Faculty of Health and Life Science, Daffodil International University, Dhaka, Bangladesh

**Keywords:** Surface electromyography, handgrip dynamometry, heart rate recovery, TRIPP framework, integrative neuromuscular training, athletic performance benchmarking, sport-specific conditioning

## Abstract

**Background and Purpose:** Badminton imposes considerable cardiovascular and musculoskeletal stress. Physiological profiling can identify modifiable injury risk factors and inform exercise-based prevention and rehabilitation. This study compared cardiovascular recovery, neuromuscular activation, and limb strength between elite and recreational male university badminton players to derive preliminary physiological benchmarks for injury risk stratification and exercise rehabilitation guidance.

**Methods:** Forty male athletes (20 elite: national/university representatives with ≥5 years of competitive experience; 20 recreational: *<*3 years of experience) completed assessments of heart rate recovery (HRR), biceps brachii surface electromyography (sEMG; SENIAM protocol), handgrip strength (JAMAR dynamometry), and maximal bodyweight squat repetitions. Independent-sample *t*-tests with Cohen’s *d* (α = 0.05) and Pearson correlations were applied.

**Results:** Elite players demonstrated significantly greater handgrip strength (49.00±6.12 vs. 39.00±5.45 kg, *p* = 0.001, *d* = 1.72) and lower-limb (LL) strength (60.35±11.29 vs. 41.75±6.72 repetitions, *p <* 0.001, *d* = 1.96). Normalized sEMG root mean square (RMS) was higher in elite athletes during flexion (11.56±4.16% vs. 7.26±5.15%, *p* = 0.004, *d* = 0.94) and extension (12.67±4.56% vs. 7.85±5.73%, *p* = 0.003, *d* = 0.94). HRR did not differ significantly between groups (*p* = 0.17, *d* = 0.43, observed power = 0.34). Elite players nonetheless showed a more favorable recovery distribution. sEMG -HRR correlations were weak and non-significant in both groups.

**Conclusions:** Elite badminton players exhibit a distinct physiological profile of greater strength and more efficient neuromuscular activation. These preliminary cross-sectional findings may support the design of exercise-based injury-prevention and rehabilitation in university badminton.

## 1 Introduction

Badminton is a physiologically demanding racket sport requiring coordinated integration of aerobic endurance, anaerobic power, and neuromuscular precision [1], [2]. Elite competition involves explosive movements exceeding 6 m lunges, 60 cm vertical jumps, directional changes *>* 5 m/s, and overhead smashes *>* 400 km/h [1], [2]. Match durations frequently exceed 60 − 90 min, demanding sustained high-intensity performance in extended rallies while maintaining technical accuracy under progressive physiological fatigue [3], [4].

Recent systematic reviews report injury incidence rates of 2.5−4.4 per 1,000 training hours, with elite players aged 15 − 18 years exhibiting peak injury rates of 2.91 − 3.52 injuries per 1,000 hours [5], [6]. LL injuries comprise 41–92% of total badminton-related injuries across published studies, with ankle (13.4–18.8%), knee (18.6–18.8%), and lower-back (10.0–12.3%) injuries predominating [5], [6]. The wide range in LL injury proportions (41–92%) reflects substantial methodological heterogeneity across the source studies, including differences in injury definitions, surveillance settings, participant age groups, competitive levels, and geographical populations. Despite this variance, the consistent predominance of LL injuries across all reviewed studies underscores the need for systematic LL-focused screening and exercise-based prevention strategies in badminton, regardless of competitive context. Anterior cruciate ligament (ACL) injuries, though less frequent (3–5% of total injuries), impose career-threatening consequences with prolonged rehabilitation periods, and approximately 70% occur through the non-contact mechanisms during deceleration and pivoting [7], [8]. Shoulder injuries affect 15−20% of competitive badminton players due to repetitive overhead strokes that generate glenohumeral joint reaction forces exceeding 1.5 times bodyweight [2], [9]. These injury patterns highlight the need for systematic screening and exercise-based prevention strategies targeting modifiable physiological and neuromuscular risk factors.

The Translating Research into Injury Prevention Practice (TRIPP) framework emphasizes six consecutive stages for effective injury prevention: (1) injury surveillance, (2) analysis of etiology and mechanism, (3) development of prevention strategies, (4) efficacy testing under ideal Conditions, (5) real-world implementation, and (6) effectiveness and scalability evaluation [10], [11]. Modifiable risk factors including neuromuscular deficits, strength imbalances, inadequate cardiovascular recovery, and biomechanical maladaptation, substantially contribute to injury sus-acceptability and represent viable targets for structured training interventions[12], [13].

Cardiovascular efficiency, particularly post-exertion recovery capacity, plays a dual role in badminton performance: sustaining high-intensity output across repeated rallies and mitigating injury risk associated with neuromuscular fatigue [3],[14], [15], [16]. HRR, defined as the rate of decline in heart rate following exercise cessation, serves as a validated non-invasive marker of parasympathetic reactivation and overall cardiovascular conditioning [17], [18]. Elite athletes typically exhibit faster parasympathetic restoration compared with less-trained counterparts [19].

The strength of the hand grip reflects integrated forearm and intrinsic hand muscle function essential for racket stabilization during high-velocity strokes [20], [21], [12]. Strong correlations between grip strength and shoulder lateral-rotator strength suggest that grip assessment can serve as a [14], [22] (INT) protocols incorporating grip strengthening, rotator cuff exercises, and scapular stabilization have reduced shoulder injury rates by 25–35% in racket-sport athletes [23], [13].

LL function underpins movement efficiency and injury resilience. Athletes with greater lower extremity strength and balanced hamstring-to-quadriceps ratios demonstrate 43 − 73% lower ACL injury rates [24], [15]. Randomized controlled trials (RCTs) confirm that 8–12 weeks of INT combining balance training, plyometric, agility, and sport-specific exercises significantly reduce lower-limb injury rates while improving performance [23], [13], [25].

Surface EMG captures motor-unit recruitment, providing quantitative indices of muscle activation amplitude, timing, and neuromuscular control [17], [20], [26]. In racket sports, elite athletes exhibit greater motor-unit synchronization, reduced antagonist co-contraction, and more efficient force production than less-skilled players [21], [27]. Neuromuscular deficits, including delayed activation, excessive co-contraction, and limb asymmetry *>*10–15%, are established risk factors for lower-limb injury [28], [29].

Despite extensive knowledge of isolated physiological and neuromuscular factors in badminton, existing studies have typically assessed cardiovascular recovery, strength, or neuromuscular activation in isolation rather than simultaneously within a single cohort and under an explicit TRIPP-aligned injury prevention framework. While Bisschoff et al [4] examined autonomic markers in elite badminton players, and Wylleman et al [5]. Reported physiological and neuromuscular responses during match simulation, neither study linked multi-domain profiling to structured injury-prevention benchmarks in a university-level athlete population.

This study aimed to characterize and compare cardiovascular recovery (HRR), neuromuscular activation (biceps brachii sEMG), and upper- and lower-limb strength between elite and recreational male university badminton players, and to explore associations between neuromuscular activation and recovery capacity.

## 2 Materials and Methods

### 2.1 Human subjects ethical approval statement

This study was approved by the Research Ethics Committee of Jashore University of Science and Technology, Bangladesh (Approval No. ERC/FBST/JUST/2025-238). All participants provided written informed consent prior to enrollment after receiving detailed information about study procedures, potential risks and benefits, and their right to withdraw at any time without penalty. Participant confidentiality was maintained throughout data collection, analysis, and reporting.

### 2.2 Participants

Forty male badminton players from Jashore University of Science and Technology voluntarily participated in this comparative cross-sectional study. The recruitment of participants was started May, 2025 and ended January, 2026. Participants were divided into two groups:

- **Elite group** (*n* = 20): At least 5 years of competitive experience; current or former university or national team representation; participation in at least three inter-university or national tournaments per year; and structured training at least five sessions per week.
- **Recreational group** (*n* = 20): Less than 3 years of playing experience; participation limited to intramural or leisure games; no competitive tournament history; and irregular training (fewer than three sessions per week).

All participants were right-handed and reported no musculoskeletal injuries, cardiovascular conditions, or medication use affecting heart rate or neuromuscular activity in the preceding six months. Exclusion criteria included recent surgery, chronic pain, neurological disorders, or cardiovascular disease.

### 2.3 Anthropometric assessment

Baseline anthropometric and body-composition measurements were obtained using a digital body-composition analyzer (Omron Karada Scan HBF-375; Omron Healthcare Co., Ltd., Kyoto, Japan) shown in Fig. 1. Assessments took place in the morning after a 12-h overnight fast. Participants followed standardized hydration instructions for 24 h prior to testing.

**Figure 1:** Anthropometric data collection procedure. Participants stood barefoot on a calibrated digital body composition analyzer in an upright posture to record various anthropometric data under standardized laboratory conditions.

Measured variables included height (cm), body mass (kg), body mass index (BMI, kg/m^2^), body fat percentage, visceral fat level, and resting metabolic rate (kcal/day). All measurements adhered to the manufacturer’s guidelines.

### 2.4 Handgrip strength assessment

Handgrip strength of the dominant (right) hand was measured using a calibrated JAMAR hydraulic hand dynamometer (Patterson Medical, Warrenville, IL, USA) set at the second handle position shown in Fig. 2(a). Testing followed the American Society of Hand Therapists recommendations. Participants stood upright with the dominant arm alongside the body, shoulder in neutral rotation, elbow fully extended, and forearm in neutral rotation.

**Figure 2:** Data collection procedures for muscular strength assessment. (a) Handgrip strength measurement using a calibrated JAMAR hydraulic hand dynamometer with the participant standing in an upright posture. (b) Lower-limb strength assessment performed using a standardized body weight squat protocol under supervised laboratory conditions.

After a brief familiarization, participants performed three maximal voluntary contractions, each separated by 30 s of rest. Strong verbal encouragement was provided to promote maximal effort. The highest value (kg) across the three trials was used for analysis. The dynamometer was calibrated before each testing session according to manufacturer’s specifications.

### 2.5 Lower-limb strength assessment

Lower-limb muscular endurance was assessed using a standardized bodyweight squat repetition test conducted under the supervision of trained exercise scientists shown in Fig. 2(b). A bodyweight squat test was selected as a practical, field-friendly proxy for LL functional endurance relevant to repeated-sprint and lunge demands in badminton; this measure is noted as an endurance index rather than a direct measure of maximal strength or power, which is acknowledged as a study limitation. Participants performed a standardized warm-up consisting of 10 bodyweight squats and dynamic leg stretches, then completed as many technically correct full squats as possible across three trials (2 min rest between trials). Squat technique criteria included: (1) feet shoulder-width apart; (2) descent until thighs were parallel to the floor (knee angle ≈ 90◦); (3) controlled tempo (1 s descent, 1 s ascent); and (4) maintenance of a neutral spine. Repetitions not meeting these criteria were excluded. The highest number of correctly executed repetitions across the three trials was used for analysis.

### 2.6 Heart rate recovery protocol

HRR was assessed in the Bir Shreshtha Hamidur Rahman Central Gymnasium and Conditioning Laboratory. Participants attended testing at a consistent time of day within a 2-h window to minimize circadian effects and avoided caffeine, alcohol, and strenuous exercise for 24 h beforehand.

Resting heart rate was measured with a Polar H10 heart-rate monitor after 5 min of seated rest. Participants then performed a standardized dynamic warm-up followed by a graded treadmill protocol beginning at 6 km/h and increasing by 0.5 km/h every 30 s until moderate exertion (approximately 120 beats per minute) was reached. The data collection steps are graphically represented in Fig. 3. On Day 1, each participant’s submaximal intensity was identified as 70% of heart rate at the first kilometer. On Day 2, participants ran at this individualized intensity for exactly 5 min. Immediately after exercise, they sat quietly while heart rate was recorded every 30 s for 5 min.

**Figure 3:** Heart rate recovery (HRR) measurement protocol. Participants completed a maximal running to elicit peak heart rate and then transitioned immediately to a standardized recovery position, during which heart rate decline was recorded to quantify post-exercise autonomic recovery.

HRR was defined as the absolute heart rate recorded at 5 min post-exercise, with lower values indicating better parasympathetic recovery. It is acknowledged that a submaximal treadmill based protocol differs from sport-specific intermittent badminton play; this standardized approach was adopted for reproducibility and cross-group comparability, and its limitations are discussed below.

### 2.7 Surface electromyography assessment

#### 2.7.1 Data acquisition

sEMG data were collected in the Biomedical Signal Processing Laboratory using a BIOPAC MP45 system (BIOPAC Systems, Inc., Goleta, CA, USA) at a sampling rate of 1,000 Hz. Bipolar surface electrodes were placed over the biceps brachii of the dominant arm according to SENIAM recommendations. Skin was shaved, abraded, and cleaned to maintain impedance below 5 kΩ. Participants sat with the elbow flexed to 90◦. They performed two elbow flexions followed by two elbow extensions at a comfortable pace, repeated three times with 30 s rest between sets.The complete process of recording sEMG signal acquisition has been shown in Fig. 4. A standardized flexion–extension task was deliberately selected over sport-specific movements to ensure high inter-session reliability and to provide a baseline measure of neuromuscular control capacity applicable across skill levels; the implications of this non-sport-specific task are discussed in the Limitations section. The cleanest trial free of visible artifacts was selected for analysis.

**Figure 4:**
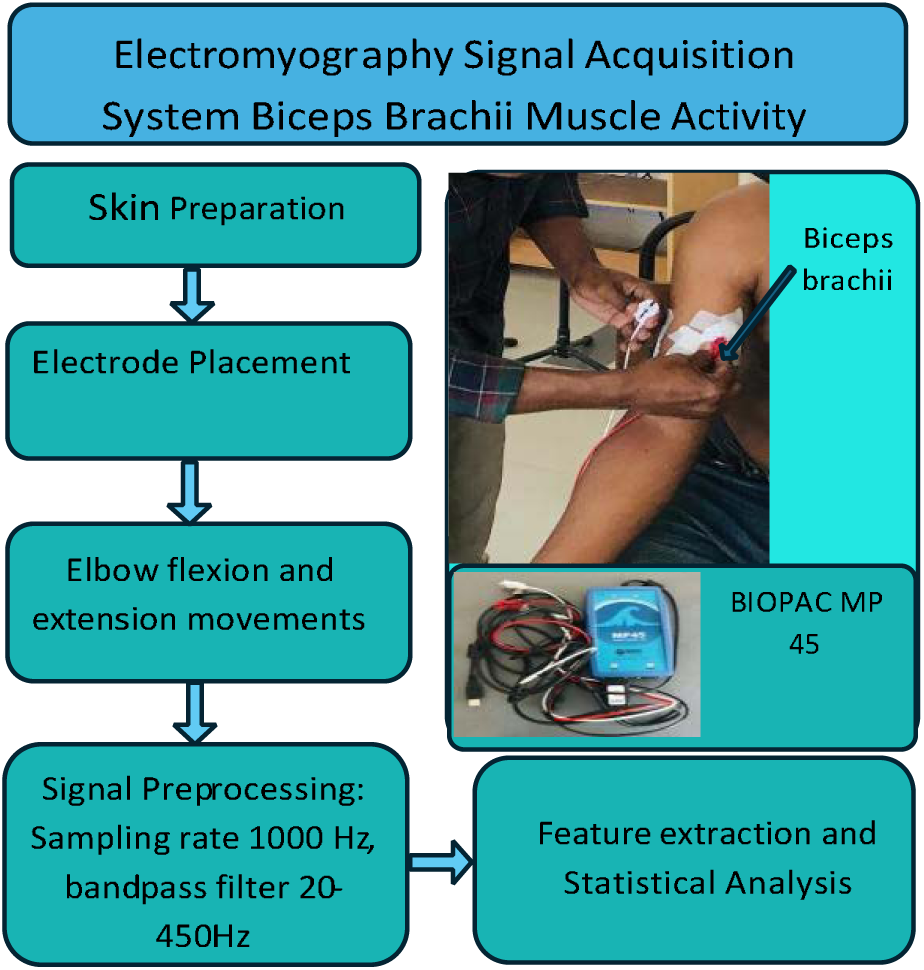
Surface electromyography (sEMG) data collection procedure. Bipolar surface electrodes were placed over the biceps brachii muscle according to standardized anatomical landmarks, and muscle activity was recorded during the elbow extension and flexion tasks.

#### 2.7.2 Signal processing and feature extraction

Data were processed in MATLAB R2023b (MathWorks, Natick, MA, USA). A fourth-order Butterworth bandpass filter (20–450 Hz) removed motion artifacts and high-frequency noise, and a 50-Hz notch filter (quality factor *Q* = 30) suppressed power-line interference. Signals were full-wave rectified to obtain amplitude envelopes. Active muscle periods were identified using a threshold-based approach: segments exceeding 10% of the maximum rectified amplitude for at least 200 ms were considered active. Root mean square (RMS) amplitude was computed using a 250-ms moving window. RMS values were normalized to the maximum RMS recorded during a maximal voluntary contraction and expressed as percent of maximum voluntary contraction (%MVC). Mean normalized RMS values for flexion and extension phases were computed for Subsequent statistical analyses.

### 2.8 Statistical analysis

A post-hoc power analysis indicated that to detect a moderate effect (d=0.5) for HRR with =0.05 and power=0.80, a sample of approximately 51 participants per group would be required. All statistical analyses were conducted in MATLAB R2023b. Descriptive statistics (mean ± standard deviation) were calculated for all variables. The Jarque–Bera test evaluated normality; all variables satisfied normality assumptions (*p >* 0.05), permitting parametric analyses.

The group differences between elite and recreational players were assessed using independent samplesVancouver press *t*-tests. The *t* statistic was computed by Eq. 1

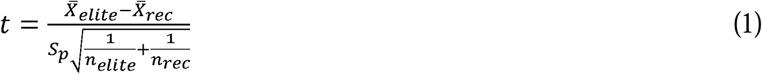

Where *X̂* denotes group means, *s_p_* is the pooled standard deviation, and *n* the group sample sizes. Statistical significance was set at α = 0.05 (two-tailed).

Effect sizes (Cohen’s *d*) were calculated by Eq. 2.

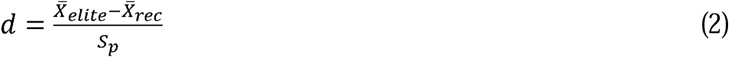

Effect-size magnitudes were interpreted as very small (*d* = 0.01), small (*d* = 0.20), medium (*d* = 0.50), large (*d* = 0.80), very large (*d* = 1.20), and huge (*d* = 2.00)[20], [26], [30]. Post-hoc observed power for each comparison was estimated in MATLAB. Pearson correlation coefficients quantified associations between neuromuscular activation (RMS) and cardiovascular recovery (HRR). Correlations were interpreted as negligible (*r <* 0.30), low (0.30–0.49), and moderate (0.50– 0.69), high (0.70–0.89), and very high (*r* ≥ 0.90)[31].

### 2.9 Use of Generative Artificial Intelligence

Generative artificial intelligence tools (e.g., large language models) were used solely to assist with language editing, sentence restructuring, and improvement of clarity in the drafting and revision of this manuscript. These tools were not used for study design, data collection, statistical analysis, or interpretation of results, and they did not generate any scientific content, numerical values, or conclusions. All scientific decisions, data handling procedures, and final manuscript content were determined, verified, and approved by the authors, who take full responsibility for the integrity and accuracy of the work.

## 3 Results

### 3.1 Anthropometric characteristics

Table 1 summarizes the anthropometric profiles. Elite players were slightly older, taller, and heavier than recreational players, but none of these differences reached statistical significance (*p >* 0.05). Body-composition metrics (BMI, body fat, visceral fat) were comparable between groups, suggesting that performance differences were not primarily attributable to gross anthropometric disparities.

**Table 1:**
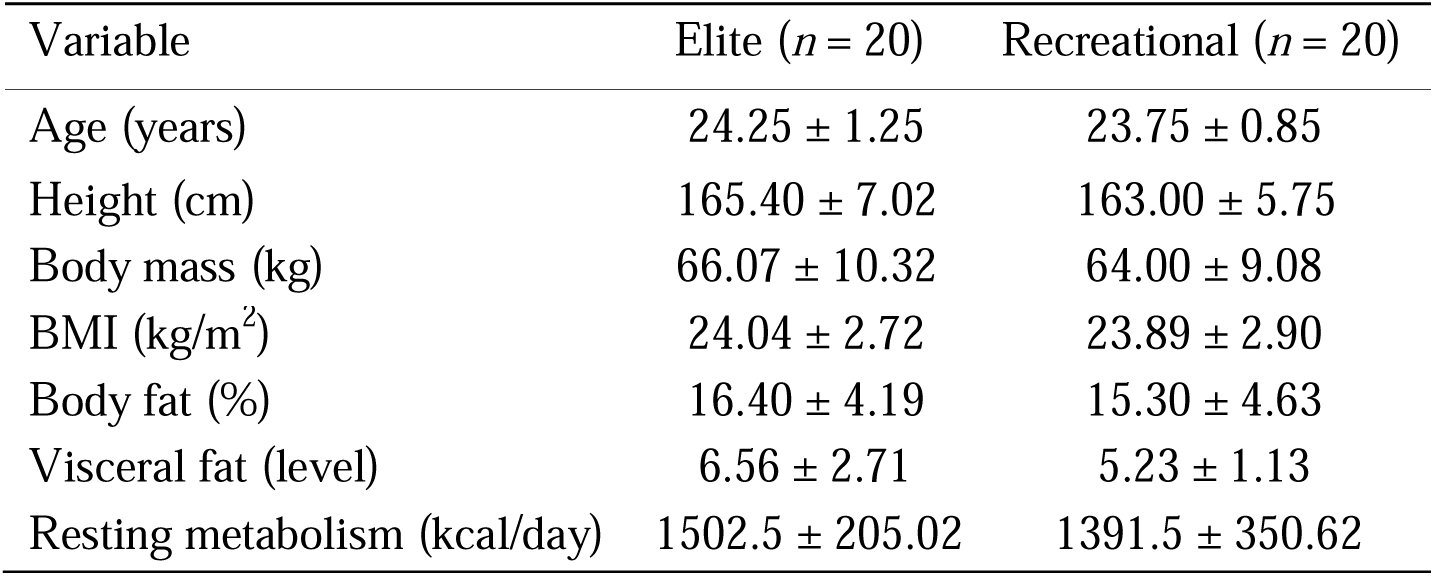
Baseline anthropometric and physiological characteristics of elite and recreational male university badminton players.

| Variable | Elite ($n = 20$) | Recreational ($n = 20$) |
| --- | --- | --- |
| Age (years) | $24.25 \pm 1.25$ | $23.75 \pm 0.85$ |
| Height (cm) | $165.40 \pm 7.02$ | $163.00 \pm 5.75$ |
| Body mass (kg) | $66.07 \pm 10.32$ | $64.00 \pm 9.08$ |
| BMI ($\text{kg}/\text{m}^2$) | $24.04 \pm 2.72$ | $23.89 \pm 2.90$ |
| Body fat (%) | $16.40 \pm 4.19$ | $15.30 \pm 4.63$ |
| Visceral fat (level) | $6.56 \pm 2.71$ | $5.23 \pm 1.13$ |
| Resting metabolism (kcal/day) | $1502.5 \pm 205.02$ | $1391.5 \pm 350.62$ |

### 3.2 Handgrip strength

Elite players demonstrated significantly greater handgrip strength than recreational players (49.00±6.12 vs. 39.00±5.45 kg; *t* (38) = 5.38, *p* = 0.001, 95% CI [6.18, 13.82], *d* = 1.72, observed power= 0.99) shown in Fig. 5. This very large effect size indicates a substantial functional advantage in upper-limb strength consistent with the demands of racket control and stroke execution in competitive badminton.

**Figure 5:**
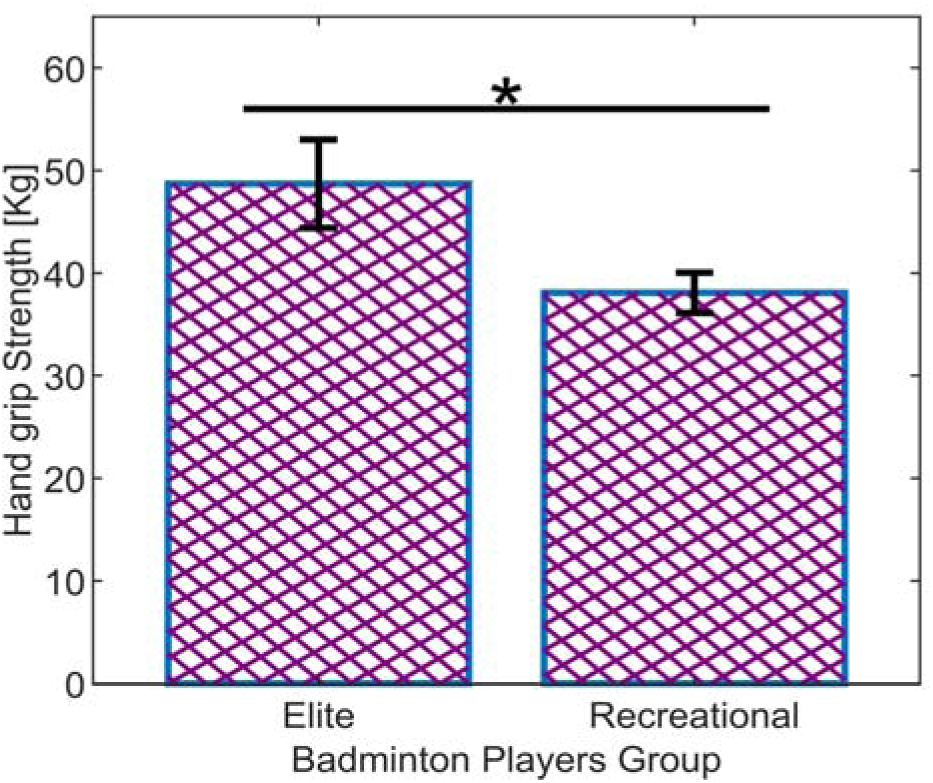
Comparison of handgrip strength between elite and recreational badminton players. Data are presented as mean ± standard deviation. Elite athletes exhibited significantly greater grip strength (*p* = 0.001), reflecting superior upper limb muscular capacity essential for racket control and stroke precision.

### 3.3 Lower-limb muscular endurance

Elite players completed significantly more maximal bodyweight squat repetitions than recreational players (60.35±11.29 vs. 41.75±6.72; *t* (38) = 6.29, *p <* 0.001, 95% CI [12.67, 24.53], *d* = 1.96, observed power= 1.00) shown in Fig. 6. This very large effect size highlights the critical role of LL muscular endurance in supporting rapid, repeated movements in high-level badminton.

**Figure 6:**
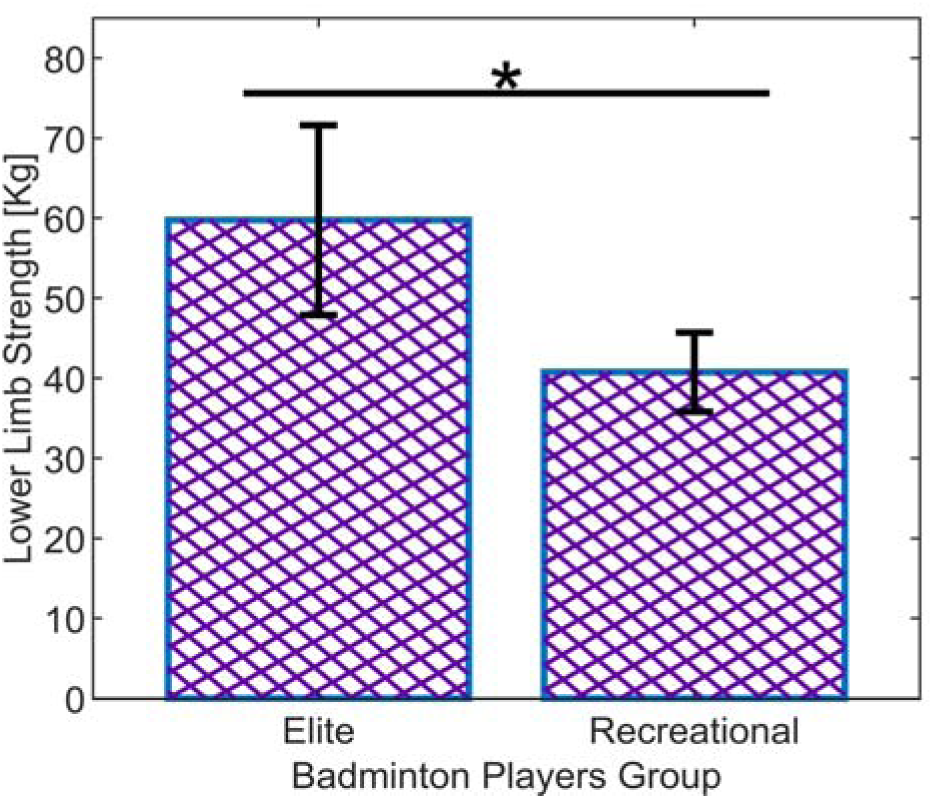
Comparison of LL strength (maximal bodyweight squat repetitions) between elite and recreational badminton players. Data are presented as mean ± standard deviation. Elite athletes performed significantly more repetitions (*p <* 0.001), indicating superior lower body muscular endurance and functional capacity.

### 3.4 Neuromuscular activation: biceps brachii sEMG

During elbow flexion, elite players exhibited significantly higher normalized RMS amplitude compared with recreational players (11.56±4.16% vs. 7.26±5.15%; *t* (38) = 2.98, *p* = 0.004, 95% CI [1.42, 7.18], *d* = 0.94, observed power= 0.87) shown in Fig. 7. This large effect suggests more efficient motor-unit recruitment and greater neuromuscular activation during concentric contractions, with lower variability indicating more consistent control.

**Figure 7:**
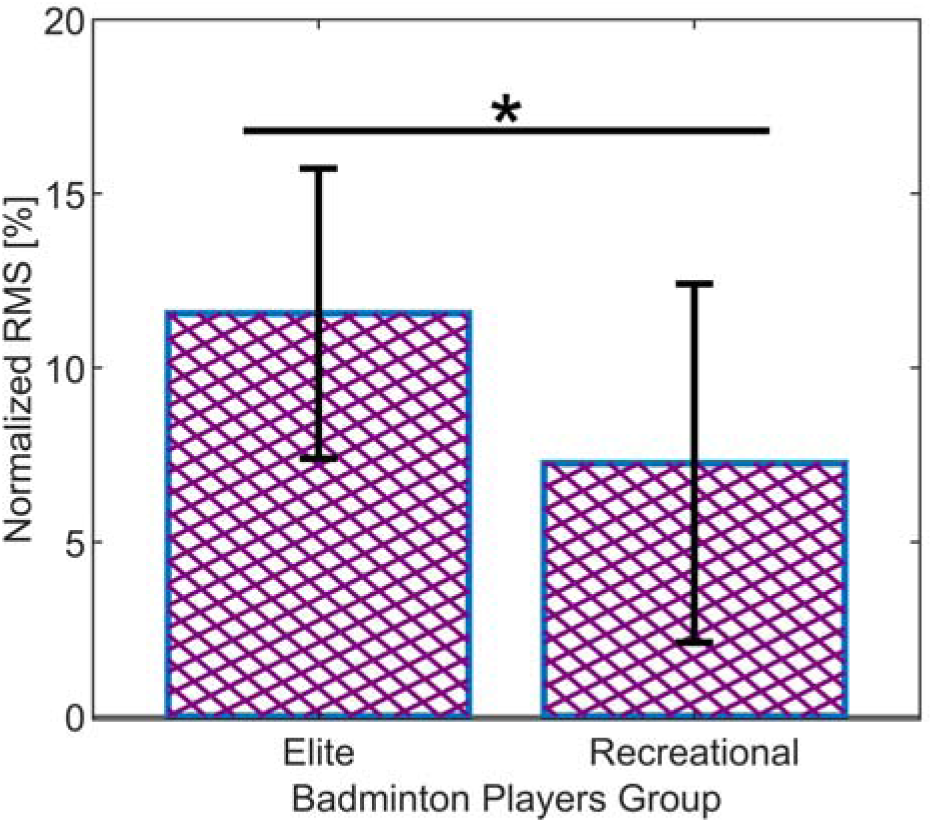
Comparison of normalized RMS amplitude of biceps brachii during elbow flexion between elite and recreational badminton players. Data are presented as mean ± standard deviation. Elite players demonstrated significantly higher neuromuscular activation (*p* = 0.004), reflecting superior motor unit recruitment and coordination.

Similarly, during the elbow extension, elite players showed higher normalized RMS amplitudes (Fig. 8) (12.67±4.56% vs. 7.85±5.73%; *t* (38) = 3.01, *p* = 0.003, 95% CI [1.77, 7.87], *d* = 0.94, observed power= 0.88), indicating enhanced neuromuscular activation even in eccentric or stabilization phases, supporting controlled deceleration and precise movement execution.

**Figure 8:**
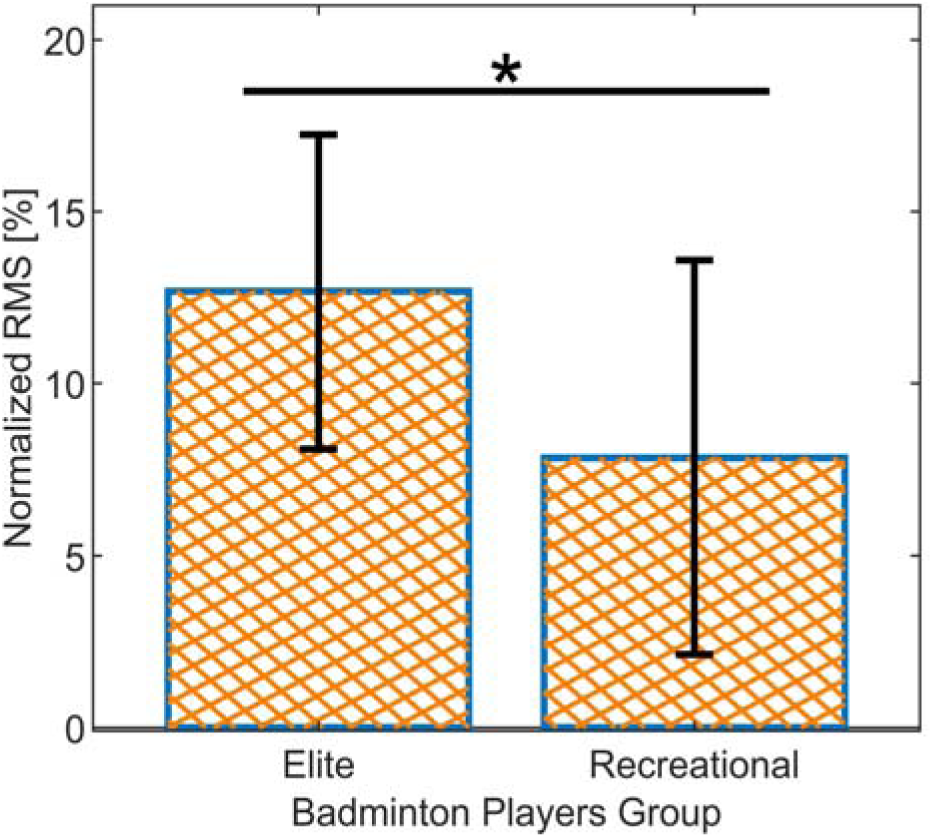
Comparison of normalized RMS amplitude of biceps brachii during elbow extension between elite and recreational badminton players. Data are presented as mean ± standard deviation. Elite players demonstrated significantly higher neuromuscular activation (*p* = 0.003), reflecting superior motor unit recruitment and coordination.

### 3.5 Heart rate recovery

Mean HRR at 5 min post-exercise was numerically higher bpm in elite players (74.60±9.70 vs. 71.15±6.05 bpm), but the difference was not statistically significant *t* (38) = 1.37, *p* = 0.17, 95% CI [−1.62, 8.52], *d* = 0.43, observed power= 0.34). Distributional inspection indicated that several elite players recovered into the 90–100 bpm range, while recreational players generally remained *>*100 bpm after 5 min, suggesting a non-significant trend toward faster autonomic recovery in the elite group (Fig. 9). This null result should be interpreted cautiously given the modest observed power (0.34), which indicates insufficient statistical power to detect moderate effects with this sample size.

**Figure 9:**
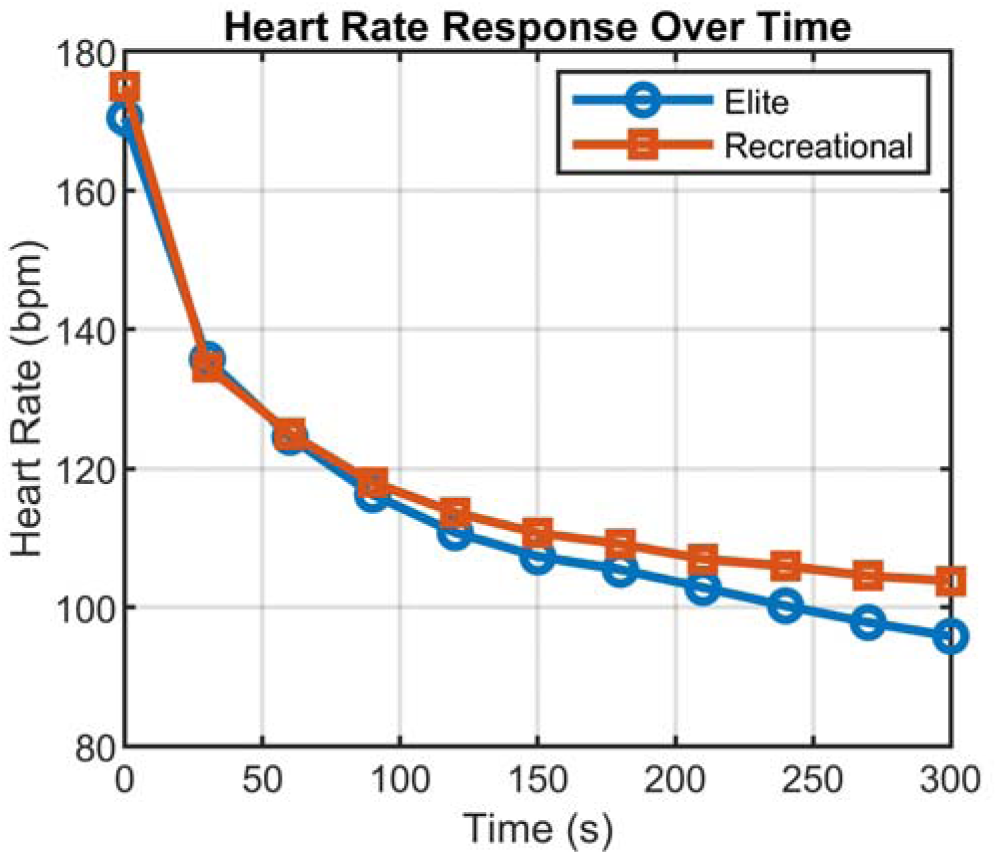
Distribution of HRR values at 5 minutes post-exercise.

### 3.6 Correlation between neuromuscular activation and cardiovascular recovery

In elite players, the correlation between mean biceps brachii RMS activation and HRR was positive but non-significant (*r* = 0.29, *p* = 0.21), as shown in Fig. 10 (left), indicating a negligible-to-low and statistically unreliable association. In recreational players, the correlation was slightly stronger but still non-significant (*r* = 0.37, *p* = 0.10), as shown in Fig. 10(right), with larger variability in both sEMG and HRR contributing to inconsistent relationships. These weak, non-significant correlations suggest that neuromuscular and cardiovascular adaptations may develop through partially independent pathways in this cohort, though the small sample limits firm

**Figure 10:**
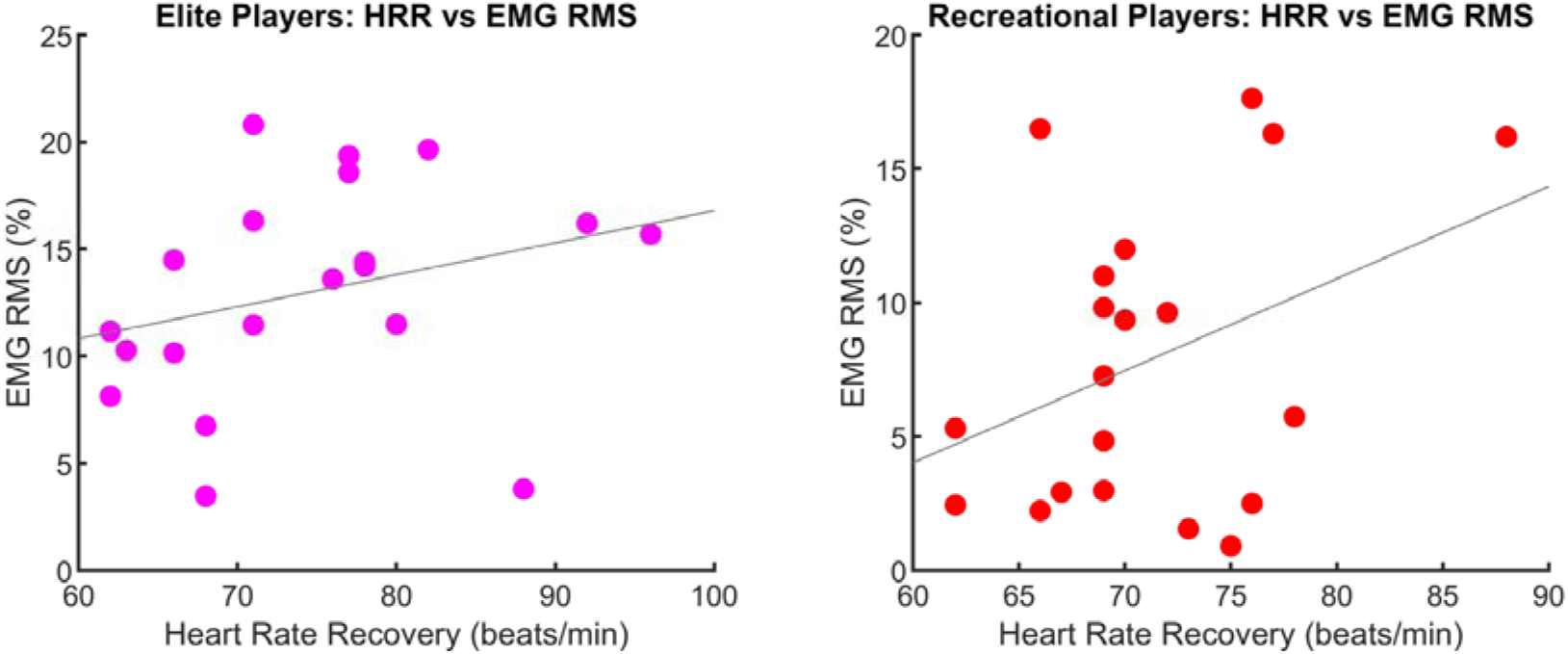
Scatter plots of normalized sEMG RMS vs. HRR in elite (left) and recreational (right) players.

Conclusions.

## 4 Discussion

This cross-sectional study compared cardiovascular recovery, neuromuscular activation, and upper- and lower-limb strength between elite and recreational male university badminton players with the aim of deriving preliminary physiological benchmarks for injury risk stratification and exercise rehabilitation guidance. With respect to the primary objective, HRR at 5 min post exercise did not differ significantly between groups (*p* = 0.17, *d* = 0.43), and observed statistical power was modest (0.34), precluding firm conclusions about autonomic differences at this sample size. Regarding the secondary objectives, elite athletes demonstrated substantially greater handgrip strength (*d* = 1.72), lower-limb muscular endurance (*d* = 1.96), and normalized biceps brachii sEMG amplitude during both flexion (*d* = 0.94) and extension (*d* = 0.94). The sEMG– HRR correlations were weak and non-significant in both groups (elite: *r* = 0.29; recreational: *r* = 0.37), consistent with the hypothesis that neuromuscular and cardiovascular adaptations may develop through partially independent pathways. These group differences, taken together, suggest a distinct physiological profile in elite players that may have implications for exercise-based injury-prevention and rehabilitation planning, though the cross-sectional design limits causal inference.

### 4.1 Handgrip strength and upper-limb injury risk

Elite players exhibited approximately 26% higher handgrip strength (*d* = 1.72), reinforcing the importance of upper-limb strength for racket control, stroke velocity, and load attenuation at the wrist and elbow. Stronger grip and forearm musculature may improve shock absorption and force distribution, which may reduce lateral epicondylitis and other overuse injuries in racket sports[32]. Epidemiological evidence confirms that rotator cuff tendinopathy and impingement remain common in competitive badminton and are linked to deficits in strength and control within the kinetic chain[33], [34]. Incorporating grip-specific and rotator cuff–focused exercises into conditioning appears to be a practical, low-cost strategy to enhance stroke performance and Support injury-preventive rehabilitation objectives[23], [13].

### 4.2 Lower-limb strength, movement efficiency, and injury prevention

Elite athletes performed about 45% more maximal squat repetitions (*d* = 1.96), indicating superior LL endurance that supports repeated accelerations, lunges, and directional changes in match play. This result is particularly relevant given that lower-extremity injuries, including strains, sprains, tendinopathies, and stress fractures that account for most badminton injuries across Levels of play[32], [5]. Evidence from RCTs shows that INT programs combining strength, balance, plyometric, and agility can improve functional movement, reduce limb asymmetry, and lower Injury risk by approximately 35–45% in racket-sport athletes[23], [13]. The large group differences in LL muscular endurance observed here support prioritizing LL resistance and plyometric training as core components of exercise-based injury-prevention and return-to-sport rehabilitation programs.

### 4.3 Neuromuscular activation, motor control, and eccentric capacity

Elite players demonstrated about 60% higher normalized biceps brachii activation during both flexion and extension (*d* = 0.94), suggesting more efficient motor-unit recruitment and greater eccentric control during arm deceleration. These findings align with EMG-guided stretch shortening-cycle training studies that report improved latency, reactive strength, and impulse Characteristics in elite racket-sport athletes[1], [35]. Eccentric strength is increasingly recognized as a key protective factor against tendon overload and muscle strain, and EMG-based biofeedback can help correct asymmetries and improve movement quality during both training and rehabilitation[36], [37]. It must be noted, however, that these sEMG data were obtained during a standardized, non-sport-specific elbow task; direct inference to badminton-specific overhead or smash mechanics would require further validation using sport-specific protocols.

### 4.4 Heart rate recovery, autonomic function, and load management

The primary hypothesis that elite players would demonstrate faster HRR kinetics was not supported at the current sample size (*p* = 0.17, *d* = 0.43, observed power = 0.34). This null result does not exclude a meaningful autonomic difference between groups. It reflects insufficient statistical power to detect a moderate effect with *n* = 40. A post-hoc power analysis indicates that detecting *d* = 0.43 at α = 0.05 with 80% power would require approximately 88 participants per group. The moderate effect size and more favorable HRR distribution in elite players nevertheless remain compatible with enhanced parasympathetic reactivation typically observed in trained athletes.

### 4.5 Integrated implications for exercise-based injury prevention and rehabilitation

The present findings indicate that elite badminton players combine greater strength, more efficient neuromuscular activation, and numerically more favorable recovery profiles. These combined differences may reflect a physiological profile associated with enhanced performance and potentially reduced injury susceptibility, though the cross-sectional design precludes causal conclusions. In applied practice, three exercise-based strategies are supported by the present data alongside broader evidence: (1) routine profiling of grip strength, LL muscular endurance, simple functional tests, and HRR to identify potentially high-risk athletes and guide exercise program individualization; (2) implementation of 8–12 week INT blocks (2–3 sessions/week) emphasizing LL strength, balance, plyometric, core stability, and shoulder/forearm conditioning to address modifiable risk factors[6], [13]; and (3) basic HRR monitoring to individualize training load and identify early signs of maladaptation or overuse risk [38].

### 4.6 Limitations and future research directions

This study has several important limitations that should be considered when interpreting findings. First, the modest sample size (*n* = 40) and cross-sectional design limit statistical power for detecting moderate effects (particularly for HRR, observed power= 0.34) and preclude causal inference about training adaptations or injury risk. Additionally, participants were recruited from a single university using a convenience sampling approach, which limits the generalizability of findings to other institutions, competitive levels, or cultural contexts. Second, the exclusively male cohort restricts generalizability to female athletes, who may exhibit distinct injury profiles and neuromuscular characteristics. Third, neuromuscular assessment was limited to biceps brachii sEMG during isolated tasks, and cardiovascular recovery was assessed after treadmill running rather than sport-specific intermittent play. Finally, the absence of prospective injury surveillance means that links between physiological markers and actual injury incidence remain Inferential.

Future research should prioritize 6–12 month longitudinal RCTs integrating INT protocols with comprehensive injury tracking to confirm causal benefits and validate injury-prevention effectiveness. Studies should expand neuromuscular profiling to include LL and shoulder musculature during badminton-specific movements (e.g., lunges, overhead strokes) and establish sex specific normative data. Development of multivariable risk-prediction models and deployment of wearable sensors for real-time on-court monitoring may further refine personalized exercise-based injury-prevention and return-to-sport strategies.

## 5 Conclusion

Elite badminton athletes in this university cohort exhibited substantially greater handgrip strength (*d* = 1.72), LL muscular endurance (*d* = 1.96), and neuromuscular activation (*d* = 0.94) than recreational players, representing a distinct physiological profile that may be associated with reduced injury susceptibility through enhanced joint stability and load distribution. HRR did not differ significantly between groups, although a moderate effect size and more favorable recovery distribution were observed in elite players. To our knowledge, this study provides the first integrated cross-sectional profile combining cardiovascular recovery, sEMG-based neuromuscular assessment, and multi-limb strength evaluation within a university badminton cohort in South Asia, contextualized within the TRIPP framework.

Based on the present cohort data, preliminary screening reference values of handgrip strength ≥ 45 kg, squat repetitions ≥ 55, and normalized sEMG RMS ≥ 10% MVC may warrant further investigation as candidate thresholds for injury risk stratification; however, these values are cohort-specific and should be validated in larger, multi-centre, prospective studies before adoption in routine screening or rehabilitation protocols. Routine physiological profiling incorporating handgrip strength, lower-limb muscular endurance, and sEMG-based neuromuscular assessment combined with structured INT implementation may provide an accessible, scalable strategy for reducing badminton’s injury burden while supporting performance excellence. The present data are preliminary and cross-sectional; confirmation by longitudinal, prospective, multi-centre studies with adequate statistical power is required before these benchmarks are adopted in routine clinical or rehabilitation screening.

## Supporting information

https://drive.google.com/file/d/1XXyuuE4XNtrZb0q4ZF_4XjHXKKz95__v/view?usp=drive_link

https://drive.google.com/file/d/1yxSZR92raiou_2LmT1C63WsuwOSthFP9/view?usp=drive_link

## Acknowledgments

We gratefully acknowledge the university’s badminton coaching staff for facilitating athlete recruitment and providing access to training facilities. We thank the Department of Biomedical Engineering for providing the BIOPAC MP45 data acquisition system and signal-processing resources. Special thanks to the athletes for their time and efforts, and to the laboratory staff for assistance with data collection and manuscript preparation. The authors also acknowledge the use of generative artificial intelligence tools (large language models) to assist with language editing, sentence restructuring, and improvement of clarity during manuscript preparation.

## DECLARATIONS

### Consent for publication

Consent for publication was obtained from all participants who contributed their data for this study.

### Data availability

The datasets generated and analyzed during the current study are available from the corresponding authors upon reasonable request, subject to ethical considerations regarding participant privacy and institutional data-management policies.

### Competing Interests

The authors have no relevant financial or non-financial interests to disclose.

### Funding

This work was supported by the University Grants Commission Bangladesh funded University Grants [24-FoET-09].

### Authors’ contribution

All authors contributed to the study conception and design. HA: Conceptualization, Methodology, Resources, Writing –original draft. MM: Conceptualization, Formal Analysis, Investigation, Methodology, Project administration, Visualization, Writing original draft. MKH: Formal Analysis, Investigation, Visualization, Writing–review & editing, Writing original draft. JAS: Formal Analysis, Writing–original draft. MH: Methodology, Supervision, Visualization, Writing–review & editing. AA: Methodology, Data curation, Writing–review & editing. FAB: Investigation, Methodology, Validation, Visualization, Writing–review & editing. All authors reviewed the manuscript.

### Ethical approval

This study was approved by the Ethical Review Committee of Jashore University of Science and Technology, Bangladesh (Approval No. ERC/FBST/JUST/2025-238). All participants provided written informed consent after receiving detailed information about the study procedures, potential risks and benefits, and their right to withdraw at any time without penalty.

Participant confidentiality was maintained throughout data collection, analysis, and reporting.

### Declaration of Generative AI and AI-Assisted Technologies in the Writing Process

During the preparation of this manuscript, the authors used large language model-based generative AI tools (e.g., Perplexity) solely to assist with language editing, sentence restructuring, and improvement of grammatical clarity and readability. These tools were not used for study design, data collection, data analysis, interpretation of results, or generation of any scientific content, numerical values, figures, tables, or conclusions. All scientific decisions, analytical procedures, and final manuscript content were determined, verified, and approved by the authors, who take full responsibility for the integrity and accuracy of the work. After using these tools, the authors reviewed and edited all AI-assisted output and take full responsibility for the content of the manuscript.

### Patient privacy and data protection

All personal identifiers were removed from the dataset before analysis and reporting. Data were stored on password-protected institutional computers with access restricted to the research team only. No individual participant can be identified in any table, figure, or text of this manuscript.

## Abbreviation List

ACL: Anterior cruciate ligament
BMI: Body mass index
HRR: Heart rate recovery
INT: Integrative neuromuscular training
LL: Lower limb
MVC: Maximal voluntary contraction
RCT: Randomized controlled trial
RMS: Root mean square
sEMG: Surface electromyography
SENIAM: Surface electromyography for the non-invasive assessment of muscles
TRIPP: Translating Research into Injury Prevention Practice

## Notes

### Competing Interest Statement

The authors have declared no competing interest.

