## Supplementary material for "Cross-Sectional Physiological and Neuromuscular Profiling of Elite and Recreational University Badminton Athletes: Preliminary Benchmarks for Exercise-Based Injury Risk Stratification": https://drive.google.com/file/d/1XXyuuE4XNtrZb0q4ZF_4XjHXKKz95__v/view?usp=drive_link

ডীন-এর কার্যালয়
জীববিজ্ঞান ও প্রযুক্তি অনুষদ
যশোর বিজ্ঞান ও প্রযুক্তি বিশ্ববিদ্যালয়
যশোর-৭৪০৮, বাংলাদেশ।
ফোন : +৮৮০২ ৪২-১৪২১৬৮

Office of the Dean
Faculty of Biological Science and Technology
Jashore University of Science and Technology
Jashore- 7408, Bangladesh

সূত্র: ERC/FBST/JUST/2025-238

তারিখ: 20/05/2025

### Certificate of the ethical approval of research proposal

This is declared that the research proposal of Dr. Md. Moznuzzaman, Associate Professor, Dept. of Electrical and Electronic Engineering, Jashore University of Science and Technology has been approved complying the ethical guidelines to work on **"Comparative Study of Physical and Physiological Characteristics between Professional and Amateur University-Level Badminton Players."** associated with human model in response to his application for ethical permission.

Chairman
Ethical Review Committee
Faculty of Biological Science and Technology
Jashore University of Science and Technology
Jashore, Bangladesh

Member Secretary
Ethical Review Committee
Faculty of Biological Science and Technology
Jashore University of Science and Technology
Jashore, Bangladesh
