## Supplementary material for "Cross-Sectional Physiological and Neuromuscular Profiling of Elite and Recreational University Badminton Athletes: Preliminary Benchmarks for Exercise-Based Injury Risk Stratification": https://drive.google.com/file/d/1yxSZR92raiou_2LmT1C63WsuwOSthFP9/view?usp=drive_link

PLOS
A catalyst for better

### Human Participants Research Checklist

#### Complete the following if your study involved human participants or human participants' data. These questions should be addressed for prospective and retrospective studies.

1. Did you obtain ethics approval for this study?

- If yes, please upload the original approval document you received from your ethics committee (file type "Other").
  - Please ensure you have provided the original ethics approval document as opposed to any extension documents
  - Where ethics approval was obtained from more than one study location, please provide approval document(s) from all of the sites.
  - If the original document is in another language, please also provide an English translation.

YES
____Uploaded ____N/A

- If either of the following applies to your submission, please explain the reasons in the space below:
  - You did not obtain ethical approval.
  - You performed an international study, but did not obtain ethical approval from all countries where data collection took place.

| Not Applicable |
|---|

Continues on page 2.

2. If you prospectively recruited human participants for the study – for example, you conducted a clinical trial, distributed questionnaires, or obtained tissues, data or samples for the purposes of this study, please report in the Methods:

   i. the day, month and year of the **start and end** of the recruitment period for this study.

   ii. whether participants provided informed consent, and if so, what type was obtained (for instance, written or verbal, and if verbal, how it was documented and witnessed). If your study included minors, state whether you obtained consent from parents or guardians. If the need for consent was waived by the ethics committee, please include this information.

Please state the line number(s) in the Methods where this is reported 171,177

YES Completed ____N/A

3. If you are reporting a retrospective study of, for example, medical records, archived samples, survey data, please report in the Methods section:

   ii. the day, month and year when the data were accessed for research purposes

   iii. whether authors had access to information that could identify individual participants during or after data collection.

Please state the line number(s) in the Methods where this is reported _______

____Completed N/A N/A
